# Synergistic Action of Commercial Lipase, Lipoxygenases and Proteinase towards PLA Depolymerization

**DOI:** 10.1101/2025.05.13.653912

**Authors:** Andrea Salini, Paolo Gonnelli, Constance Padoan, Yosra Helali, Jehan Waeytens, Salvatore Fusco, David Cannella

## Abstract

Polylactic acid (PLA) is a biobased aliphatic polyester produced from renewable resources. Its building block is lactic acid (LA), which is mainly produced through fermentation processes rather than chemical synthesis, as fermentation enables the production of optically pure D-LA or L-LA, instead of a racemic mixture. In this work, three commercial enzymes Proteinase K (PrK) from Tritirachium album, Candida rugosa lipase (CrLip), and Glycine max lipoxygenase (LOX I-B) were investigated for their synergistic action for the degradation of solvent-cast as well as EoL-PLA films. While Lip and LOX I-B alone did not promote PLA hydrolysis, they significantly enhanced PrK-mediated hydrolysis by 5.7-fold and 2.5-fold, respectively. Lip, and to a lesser extent LOX I-B, increased PrKs productive binding to the polymer, thus accelerating catalysis. These findings underline the potential of synergistic enzymes for PLA degradation and provide the first evidence of lipoxygenase activity in this process.

## Introduction

Polylactic acid (PLA) is a biobased aliphatic polyester produced from renewable resources. Its building block is lactic acid (LA), which is mainly produced through fermentation processes rather than chemical synthesis, as fermentation enables the production of optically pure D-LA or L-LA, instead of a racemic mixture [1]. Additionally, LA fermentation has the potential to utilise as feedstock non-food sources, such as lignocellulosic biomasses, reducing competition with food supplies [2], such as sugars and corn. LA produced via fermentation can be polymerised either through the polycondensation of the LA monomer or by ring-opening polymerisation (ROP) of its cyclic dimer, lactide, to form poly-L-LA (PLLA), poly-D-LA (PDLA), or poly-D,L-LA (PDLLA). While polycondensation is less expensive to operate, it generates water as a byproduct, which limits the final molecular weight of the polymer. Consequently, ROP is the preferred industrial method, as it allows the production of solvent-free polymers with higher molecular weights. However, the additional steps required for lactide synthesis and purification result in higher production costs for PLA compared to petroleum-based plastics [3]. ROP also enables the controlled incorporation of D- and L-LA monomers, facilitating the tuning of polymer properties. For instance, the inclusion of a small percentage of D-LA can affect the melting temperature (*T*_*m*_) and glass transition temperature (*T*_*g*_), improving the polymer’s performance during melt-processing conversion [4].

PLA is the most widely produced bioplastic, with a global production volume of approximately 1 million tons in 2024, which is projected to increase 2.6-fold by 2029 [5]. PLA is utilised across various sectors, including packaging, textiles, and biomedical applications, owing to its biocompatibility and the non-toxic nature of its degradation products. However, its applicability is limited by brittleness, low thermal stability, hydrophobicity, and its low rate of biodegradation in vivo [6]. In fact, PLA requires high temperatures, close to its *T*_*g*_ (i.e., 55–60 °C), to be efficiently depolymerised. In marine and terrestrial ecosystems, as well as in urban composting facilities where temperatures remain lower than PLA’s *T*_*g*_, its degradation can take decades to complete. Only under controlled conditions of industrial composting facilities, where temperatures exceed 60 °C, PLA can be completely broken down within 180 days [7]. Given PLA’s slow degradation rate in the environment and its high production cost, implementing recycling and upcycling strategies is recommended as a more sustainable approach for managing end-of-life (EoL) PLA products [8]. To do so, over the past two decades, significant research efforts have been dedicated to the discovery and characterisation of EoL-PLA degrading enzymes.

PLA depolymerases belong to the serine hydrolase class and include proteases (EC 3.4.-), which specifically target PLLA [9], as well as carboxylesterases, lipases and cutinases (EC 3.1.1.-), which preferentially hydrolyse PDLLA. PLA-active proteases are promiscuous enzymes capable of degrading PLLA due to its structural similarity to silk fibroin, which consists of L-alanine, i.e., a structural analogue to L-LA [10]. Williams (1981) [11] was the first to report enzymatic degradation of PLA, identifying Proteinase K from *Tritirachium album* as an effective enzyme, which continues to serve as a gold standard for PLA degradation research. Among commercially available enzymes, Oda et al. (2001) [12] screened 56 proteases and identified alkaline proteases as the most effective in hydrolysing PLA, linking their activity to keratin-degrading properties. Additionally, the search for PLA depolymerases has extended beyond commercial enzymes, focusing on the isolation of microbial enzymes from culture supernatants [13] and metagenomic mining approaches [14] (refs Salini et al.,). Fewer commercial lipases [15, 16] have been characterised for PLA hydrolysis and to the best of our knowledge, no studies have investigated the synergistic interactions between lipases and proteases in the context of PLA degradation.

In this work, three commercial enzymes Proteinase K (PrK) from *Tritirachium album, Candida rugosa* lipase (CrLip), and *Glycine max* lipoxygenase (LOX I-B) were investigated for their synergistic action for the degradation of solvent-cast as well as EoL-PLA films. While Lip and LOX I-B alone did not promote PLA hydrolysis, they significantly enhanced PrK-mediated hydrolysis by 5.7-fold and 2.5-fold, respectively. Lip, and to a lesser extent LOX I-B, increased PrK’s productive binding to the polymer, thus accelerating catalysis. These findings underline the potential of synergistic enzymes for PLA degradation and provide the first evidence of lipoxygenase activity in this process. This approach provides a solid foundation for the development of more efficient enzymatic cocktails for recycling and upcycling of PLA, offering sustainable alternatives for the valorisation of plastic waste.

## Materials and Methods

### 5.1.1 Materials and enzymes

All reagents used in this study were purchased from Carl Roth (Karlsruhe, Germany) and were of analytical grade unless otherwise specified. Virgin PLA (_V_PLA) films were prepared as described in section 5.1.2 using Luminy® LX175 PLA pellets (Corbion). Post-consumer PLA (_PC_PLA) was obtained from surgical mask packaging wrap. The enzymes used in this study included lipase from *Candida rugosa* Type VII (CAS: 9001-62-1) and type I-B lipoxidase from *Glycine max* (CAS: 9029-60-1), both purchased from Sigma-Aldrich (St. Louis, MO, USA). Proteinase K (CAS: 39450-01-6) was obtained from Carl Roth.

### 5.1.2 PLA film production

PLA films were prepared using the solvent-casting method, following the protocol described by Decorosi et al. (2019) [17] with minor modifications. Briefly, 2 g of PLA pellets were dissolved in 50 mL of dichloromethane under continuous stirring until complete dissolution (approximately 30 minutes). Subsequently, 25 mL of the resulting solution was transferred into a polypropylene container (10.5 cm × 16 cm), and the solvent was allowed to evaporate at room temperature overnight.

### 5.1.3 Enzymatic degradation of PLA by commercial enzymes

Three commercial enzymes, the Proteinase K (PrK) from *Tritirachium album*, a lipase from *Candida rugosa* (CrLip), and a lipoxygenase from *Glycine max* (LOX I-B), were evaluated for their ability to degrade _V_PLA and _PC_PLA films, which were cut into 0.5 cm × 0.5 cm square pieces. All reactions were conducted in a 100 mM sodium phosphate buffer (pH 7.2 ± 0.2) containing 1% (*w/v*) PLA as the substrate. The effect of calcium (Ca^2+^), a known activating cation, on PLA degradation was assessed by supplementing the reaction mixture with CaCl□at a final concentration of 0.1 mM.

Lyophilised enzymes were dissolved in the reaction buffer to prepare 100× concentrated stock solutions. Reactions were initiated by adding the enzymes to achieve final concentrations of 0.43 mg/mL for PrK, 30 µg/mL for Lip, and either 36 µg/mL or 0.15 µg/mL for LOX I-B. The reaction mixtures were incubated at 50 °C with constant shaking at 750 rpm for up to 72 hours. Reactions were stopped by centrifugation at 10,000 × g for 5 minutes. The supernatants were subsequently filtered using 0.22 µm polytetrafluoroethylene (PTFE) membranes and analysed via HPLC. To remove oxygen (O_2_), the reaction mixtures were boiled at 90 °C for 10 minutes and cooled before enzyme addition, and the headspace of the reaction tubes was flushed with carbon dioxide (CO□) during enzyme stock addition.

D-LA was quantified using the D-lactic acid assay kit (K-DATE, Megazyme), following the manufacturer’s instructions. Total LA released during the reactions was quantified using the UltiMate 3000 UHPLC system (Thermo Fisher Scientific, Waltham, MA, USA), equipped with a photodiode array detector set at 210 nm. Chromatographic separation was performed at 65 °C using a Rezex ROA-Organic Acid H□column (Phenomenex, Torrance, CA, USA) under isocratic conditions, using 5 mM H□SO□as the mobile phase at a flow rate of 0.6 mL/min. A calibration curve was prepared with LA standards solutions ranging from 8 g/L to 0.125 g/L.

Results were expressed as the percentage of film degradation, calculated by dividing the amount of LA released during the reaction by the maximum theoretical yield of LA (LA_MAX_), as shown in Equation 1.

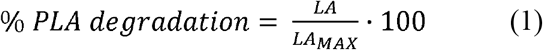

LA_MAX_ was calculated for each reaction as the total amount of LA released, assuming complete hydrolysis of the film. The value was normalised to account for the addition of a water molecule during the hydrolysis of the ester bond, which leads to an increase in the molecular weight of the monomer released compared to the monomeric unit within the polymer, as described in Equation 2.

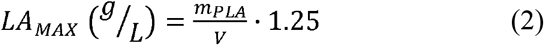

### 5.1.4 Adsorption assay

The adsorption of enzymes to PLA surface was monitored during the reaction by measuring the concentration of unbound protein in the supernatant. To achieve this, multiple reactions were prepared following the same experimental conditions described in the previous section. Aliquots of the reaction mixture were collected at the start and at the following time points (hourly in the rage 1-6 h, and at 20 h) to assess the free enzyme concentration. The protein concentration in the supernatant was quantified using the Bradford reagent (Bio-Rad, California, USA), according to the manufacturer’s instructions.

### 5.1.5 Pyruvic acid detection and quantification

High-Resolution Mass Spectrometry (HRMS) was used for targeted qualitative detection of pyruvic acid (PA) in the supernatant of the PLA incubations with enzymes. For this purpose HRMS data was acquired over the mass range of *m/z* 50–1000 using a Agilent 6546 LC/Q-TOF operated in negative mode.

HPAEC-PAD was used for quantification of PA. The separation was achieved with CarboPac PA1 column (4mm diameter x 250mm length) while the eluent program was run accordingly (Cannella et al, 2023 https://link.springer.com/article/10.1007/s10570-023-05271-z). The detection was acquired with pulsed amperometric sensor (PAD). A calibration curve was prepared with PA standard solutions ranging from 250 mg/L to 1 mg/L.

### 5.1.6 Statistical analysis

All data were collected at least in triplicate and are presented as the mean ± standard deviation. A two-tailed, completely randomised two-way ANOVA was performed to assess the statistical significance of synergistic effects within the datasets. The significance level was set at 0.05 (*), with ** indicating *p* value (*p*) ≤ 0.01; *** for *p* ≤ 0.001, and **** for *p* ≤ 0.0001.

## Results

### 5.2.1 Proteinase K and Candida Rugosa Lipase show synergism in PLA depolymerization

The PLA-depolymerising activity of Proteinase K (PrK) from *Tritirachium album* and *Candida rugosa* lipase (CrLip) was tested on solvent-cast (_V_PLA) and post-consumer PLA (_PC_PLA) films. PLA degradation is enhanced at temperatures close to its *T*_*g*_ due to the increased chain mobility, which facilitates enzymatic hydrolysis of the ester bonds. Accordingly, the incubation temperature was set at 50 °C, which is close to the *T*_*g*_ of PLA (55-60 °C) and within the previously reported optimal temperature ranges of PrK [18] and Lip [19]. Both enzymes exhibited activity in neutral and alkaline pH ranges. Therefore, assays were performed at a pH of 7.2 to minimise the contribution of base-catalysed hydrolytic side reactions that occur at alkaline pH values [20].

As shown in Figure 1A, PrK exhibited comparable yet modest activity on the two PLA substrates tested, yielding a conversion to LA of 5.7% for _V_PLA and 6.9% for _PC_PLA following a 24-hour incubation timespan. In contrast, LA release was not observed when Lip was incubated with either of the PLA substrates, which is consistent with the findings reported by Hegyesi et al., (2019) [21]. Nonetheless, despite its inability to release LA when acting as a standalone enzyme, Lip exerted a significant synergistic effect when combined with PrK, enhancing the release of LA by 5.7-fold for _V_PLA and 2.7-fold for _PC_PLA.

**Figure 1.**
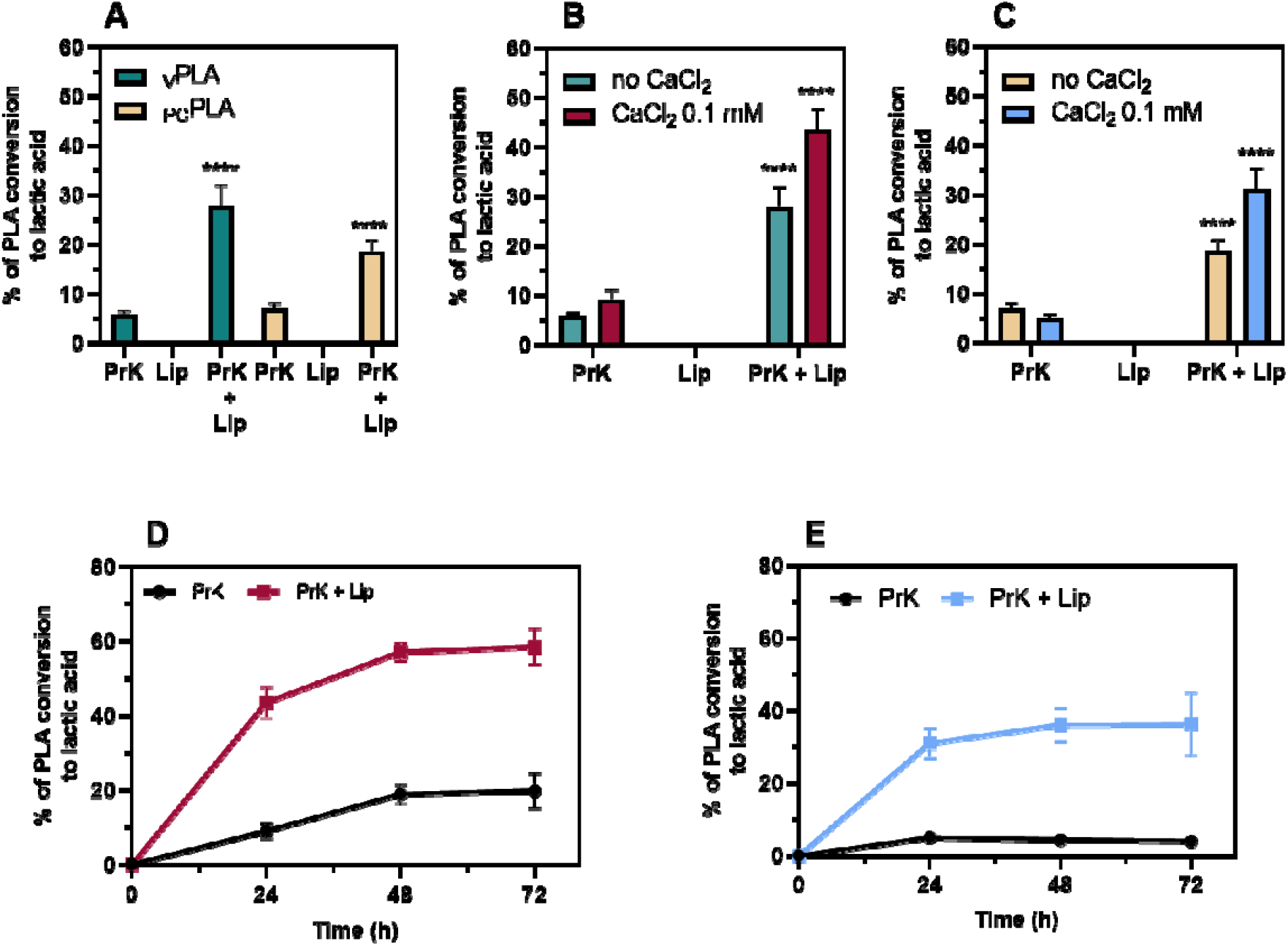
PLA film hydrolysis mediated by PrK and Lip. A) Synergistic effect of Lip in combination with PrK on _V_PLA and _PC_PLA. B) The effect of calcium on the PrK- and Lip-mediated degradation of _V_PLA. C) The effect of calcium on the PrK- and Lip-mediated degradation of _PC_PLA. D) Synergistic effect of Lip in combination with PrK on _V_PLA over 72 hours of incubation with 0.1 mM CaCl_2_. E) Synergistic effect of Lip in combination with PrK on _PC_PLA over 72 hours of incubation with 0.1 mM CaCl_2_. Enzyme concentrations used: PrK 0.43 mg/mL and Lip 30 µg/mL. Every data point was collected at least in triplicate and synergism was tested using randomised two-way ANOVA.

PrK activity is influenced by calcium, which, although not directly involved in catalysis, affects the enzyme’s three-dimensional structure. Its absence triggers a concerted conformational change which results in reduced catalytic activity [22]. To address this, the effect of adding Ca^2^□ions in the form of 0.1 mM CaCl□was tested (Fig. 1B). Interestingly, the addition of CaCl_2_ resulted in a 36% increase in activity of PrK towards _V_PLA both in the presence and absence of Lip (Figure 1B). Furthermore, Lip continued to exhibit a significant synergistic effect when combined with PrK in the presence of CaCl_2_, achieving a total conversion of 43.5% of _V_PLA to LA within a 24-hour incubation period. The extension of the reaction period led to a plateau in the combined activity of PrK and Lip, achieving approximately 60% substrate degradation in 48 hours (Figure 1D) and representing a threefold increase compared to the activity observed with PrK alone.

In the case of _PC_PLA, the addition of CaCl_2_ resulted in a 70% increase in PrK’s degradation efficiency in combination with Lip, yielding a final conversion of 31% (Figure 1C). Like the pattern observed with _V_PLA, degradation levelled off after 48 hours, leading to a total substrate hydrolysis of 36% (Figure 1E). Notably, Lip enhanced significantly by 9-fold the release of LA from _PC_PLA at 48 hours of incubation. However, a lower overall hydrolysis rate was observed compared to _V_PLA, which can likely be attributed to its higher crystallinity, which limits substrate accessibility to the enzyme.

Lipases are known to preferentially degrade PDLA [23], while PrK exhibits hydrolytic activity exclusively toward PLLA [24]. To investigate whether the synergistic effect of Lip could be attributed to its ability to target and degrade portions of the substrate composed of the D-stereoisomer, the release of D-LA was quantified (Table S1). The latter revealed that only 2–5% of the total monomer released was in the D-conformation in every sample, indicating that the synergistic action of Lip is not driven by selective degradation of D-LA-containing regions of the substrate.

### 5.2.2 Synergistic Role of LOX I-B in PrK-Mediated PLA Degradation

A similar approach was adopted to investigate the effect of *Glycine max* lipoxygenase (LOX I-B) and its potential synergistic interaction with PrK in the hydrolysis of _V_PLA and _PC_PLA. Oxidative reactions of PLA were conducted under the same experimental conditions described in the previous section, in the presence of calcium to enhance the hydrolysis yield of PrK. LOX I-B is an iron-containing redox enzyme that catalyses the oxidation of unsaturated fatty acids, to yield hydroperoxides. It has a well-established role in lipid metabolism but has never been investigated for its potential involvement in plastic oxidative degradation.

As presented in Figure 2A, LOX I-B alone showed no measurable activity toward either _V_PLA or _PC_PLA, similarly to what was previously presented for Lip. However, when combined with PrK, LOX I-B significantly enhanced substrate degradation in a concentration-dependent manner. Specifically, a higher synergistic effect was observed on _V_PLA, where LA release increased by 70% at lower LOX I-B concentrations and by 2.5-fold at higher concentrations. Although the effect was less pronounced for _PC_PLA, it was still significant, with LA release increasing by 30% and 57% at low and high enzyme dosages, respectively.

**Figure 2.**
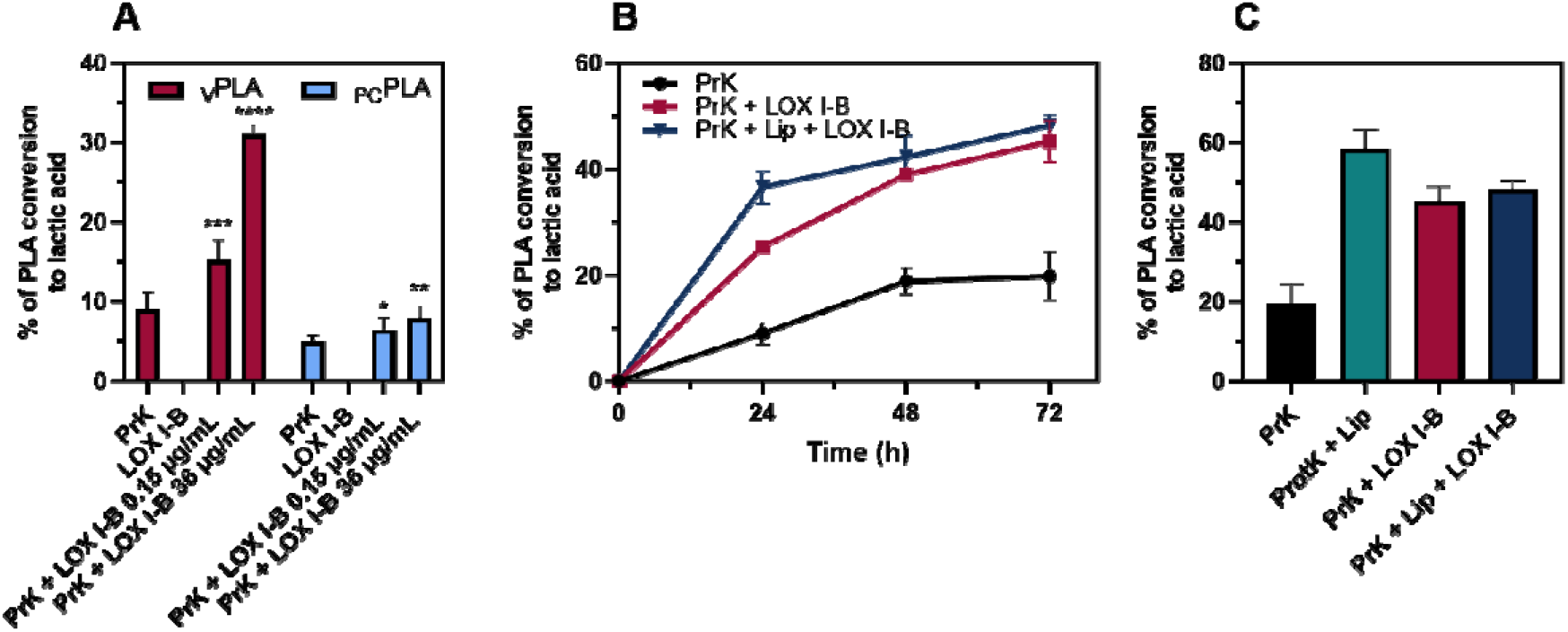
PLA film hydrolysis mediated by PrK and LOX I-B. A) Synergistic effect of LOX I-B in combination with PrK on _V_PLA and _PC_PLA. B) Time course reactions of combinations of PrK with LOX and LIP, showing the PLA conversion yield into LA. C) Specific conversion yield obtained at 72 hours for all the tested combinations of the enzymes. All reactions were carried out in 100 mM sodium phosphate buffer at pH 7.2, supplemented with 0.1 mM CaCl□and 1% (w/v) PLA film substrate. Enzyme concentrations were as follows: PrK at 0.43 mg/mL, and LOX I-B at 0.15 µg/mL and 36 µg/mL. All data was collected in triplicate and synergism was tested using randomised two-way ANOVA.

Regarding the kinetics or time course experiments presented in figure 2B, it is possible to appreciate a typical conversion kinetics for insoluble biopolymers, similar to lignocellulose derived fibres. The contribution of LOX I-B in terms of lactic acid release by PrK from PLLA remain constant for the entire course of the hydrolysis eventually reaching similar conversions obtained with a cocktail of PrK, Lipase, and Lipoxygenase after 72 hours (figure 3C). Although the highest level of PLLA conversions into LA were those obtained by the mix of lipase with PrK that yield values close to 60% of lactic acid production in 72 hours.

**Figure 3.**
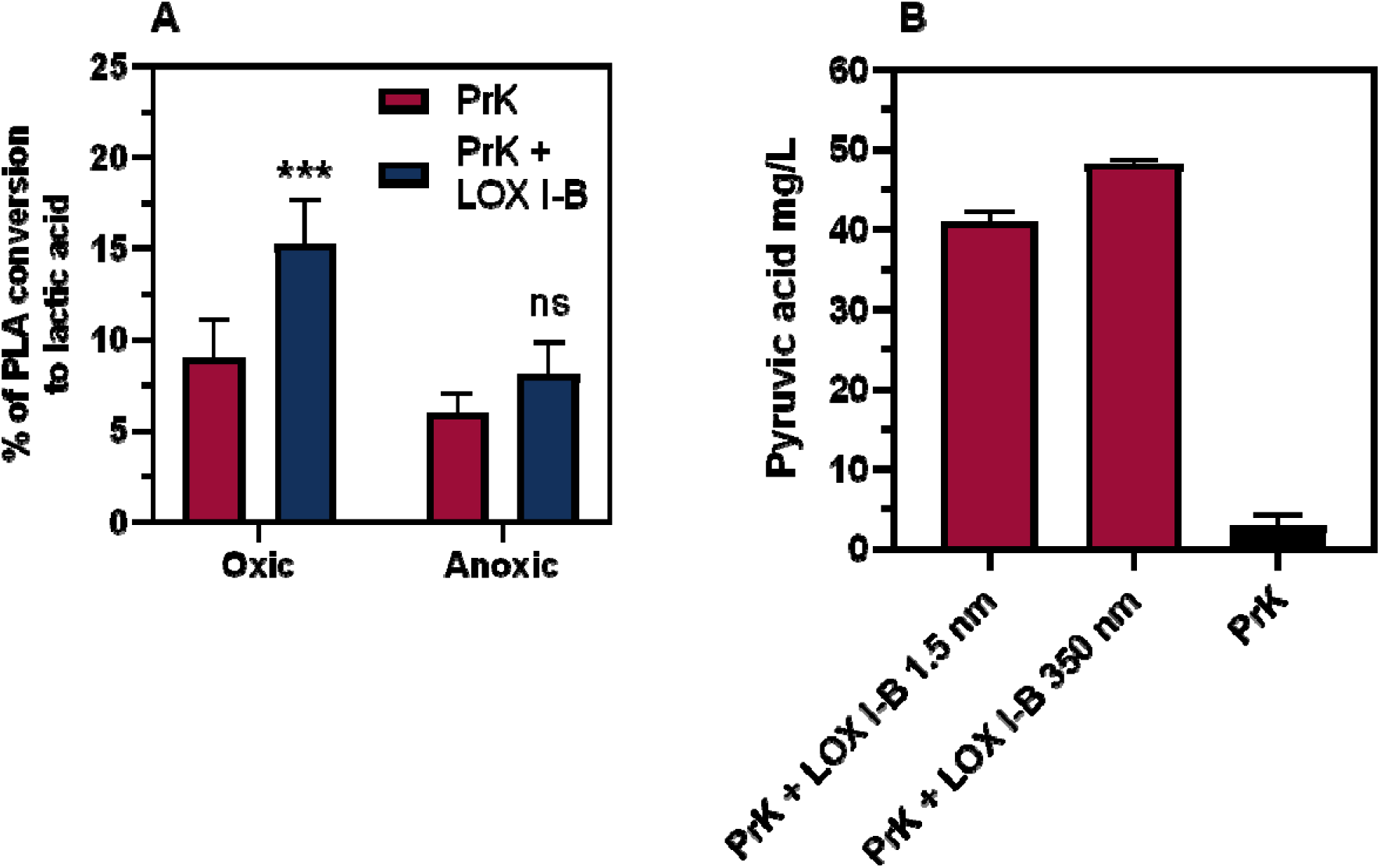
The effect of oxygen and lipoxygenase on PLA conversion to LA. A) Synergistic effect of LOX I-B in combination with PrK on _V_PLA tested in oxic and anoxic conditions. B) The quantification of pyruvic acid by HPAEC chromatography. All reactions were carried out in 100 mM sodium phosphate buffer at pH 7.2, supplemented with 0.1 mM CaCl□and 1% (w/v) PLA film substrate. Enzyme concentrations were as follows: PrK at 0.43 mg/mL, and LOX I-B at 0.15 µg/mL and 36 µg/mL. All data was collected in triplicate and synergism was tested using randomised two-way ANOVA.

**Figure 4.**
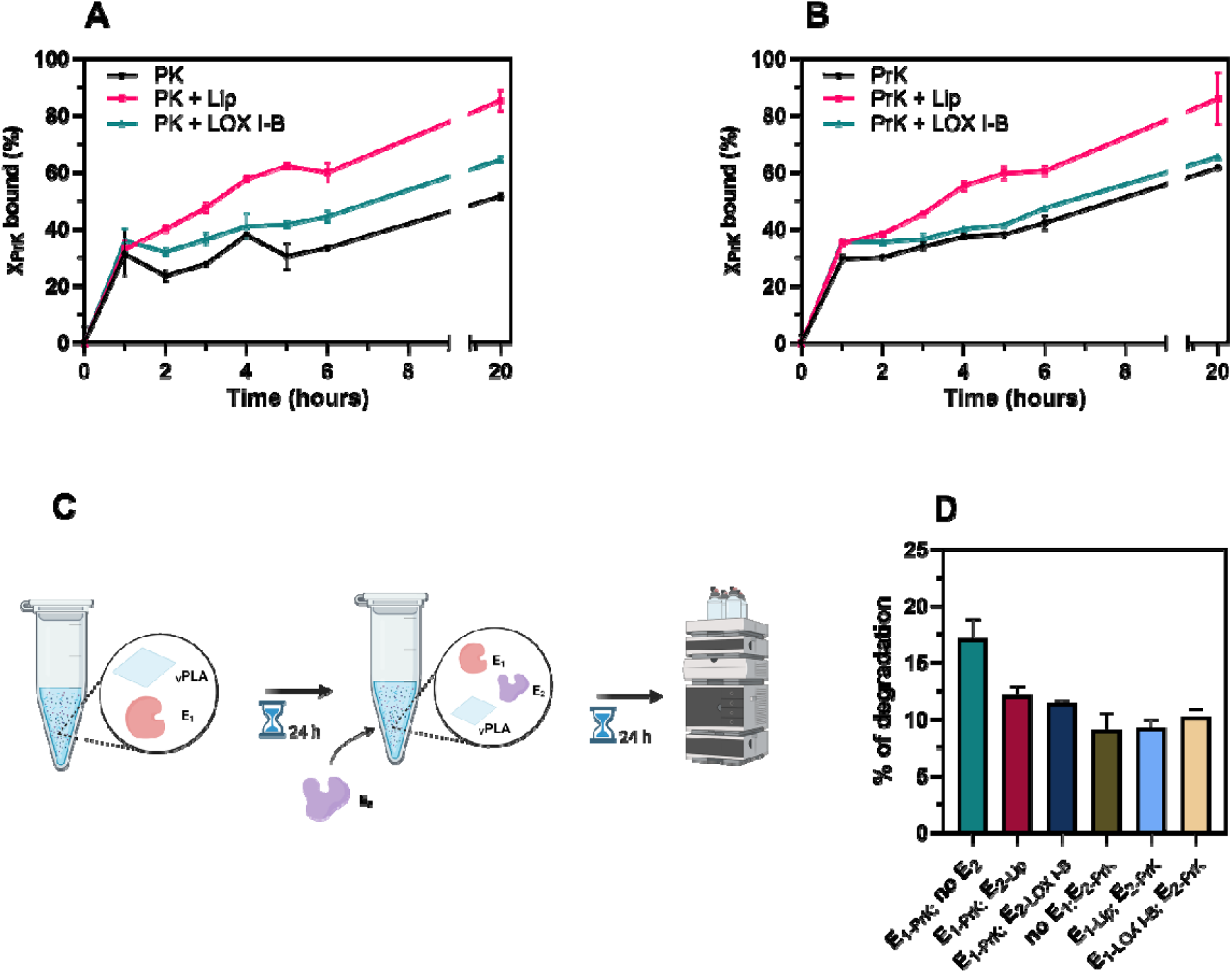
Adsorption assay and sequential addition of PrK, Lip, and LOX I-B. A, B) PrK molar fraction (χ_PrK_) bound to the substrate as a function of time during incubation on _V_PLA (A) and _PC_PLA (B), measured in the presence of Lip, LOX I-B, or PrK alone. C) Schematic representation of the reaction setup for the sequential addition of the three enzymes, as in graph D. (D) Percentage of _V_PLA conversion into lactic acid fir the sequential addition of PrK, Lip, and LOX I-B experiment: E_1_ represents the first enzyme added for 24 hours, and E_2_ represents the addition of the second enzyme after 24 hours.

These degrees of synergism towards the two different kinds of PLA suggest an interaction of LOX I-B directly on the plastic polymer structure rather than at the level of the soluble LA monomer, so ruling out a possible beneficial effect by the removal of possible product inhibition towards PrK.

### 5.2.3 LOX I-B oxidizes PLA

Since LOX I-B functions as a dioxygenase, relying on molecular oxygen (O_2_) to catalyse the oxidation of its substrates, the _V_PLA degradation assay was repeated under anaerobic conditions to assess whether its enzymatic activity was indeed responsible for the observed synergistic effect. As shown in Figure 3A, the synergistic effect exerted by LOX I-B at the lowest dosage (0,15 µg/ml), decreased significantly by 53% under anoxic conditions, resulting in degradation levels comparable to those achieved by PrK alone. The loss of synergistic action underscores the enzyme’s dependence on O_2_ to enhance PrK-mediated _V_PLA depolymerization which suggests a catalytic activity of LOX I-B on the polymer.

Two HPLC methods based on organic acid separation column (ROA H+) coupled with UV detection, as well as anion exchange chromatography (HPAEC equipped with carbopac PA1 column) coupled with amperometric PAD detector, and HRMS spectrometry were used to identify possible oxidized products from PLA upon the catalysis of LOX I-B. A consistent presence of pyruvic acid (PA) was detected in the samples from incubation of _V_PLA with LOX I-B in combination with PrK, respectively 40.9 and 48.2 mg/l (respectively for the low and high dosage tested), corresponding to 0.3 and 0.4 % of total PLA, Figure 3B. The incubation of the sole PrK on _V_PLA instead resulted into 3 mg/L of PA, close to the detection limit of the HAPAEC. No PA was detected for oxygenic incubation of LOX I-B alone with PLLA.

Moreover, to elucidate further other possible presence of lactate oxidases enzymes in the commercial LOX preparation it was also tested the direct oxidation of lactic acid (LA) into PA, either in presence or absence of _V_PLA. A lack of variation in the amount of LA before and after incubation with LOX I-B was observed, with the amount of LA that remained constant at 1 g/L as initially added, and no PA could be detected. Furthermore, lactate oxidases are known to produce hydrogen peroxide (H_2_O_2_) as a co-product of the oxidative reaction of lactate to pyruvate, so the production of H_2_O_2_ was monitored using the Amplex-red method, which allows detection to the µM scale. Also, in this case the amount of H_2_O_2_ produced was below that of the control experiments lacking the LOX I-B preparation. Overall, the data presented here does not help in elucidating a direct activity of the LOX I-B preparation towards the PLLA polymer versus a possible activity on PLA-oligomers derived from the simultaneous hydrolytic activity of PrK on PLLA.

### 5.2.4 Lipase and Lipoxygenases, promote Proteinase K adsorption to the PLLA substrate

The enzymatic hydrolysis of polyester polymers occurs in three stages: complexation, catalysis, and dissociation. The plastic film represents a large surface area to which the enzyme can adsorb. Once bound, the enzyme can displace a polymer chain and catalyse the reaction upon formation of a productive enzyme-substrate complex. After the catalysis, the enzyme and product are released from the substrate, enabling the initiation of a new catalytic cycle [25]. To assess whether Lip and LOX I-B enhanced PrK-mediated PLA hydrolysis by increasing the number of productive binding sites (BS) on the polymer, the free protein concentration was monitored during the reaction for PrK alone and in combination with Lip or LOX I-B, either on _V_PLA (Figure 3A) or _PC_PLA (Figure 3B).

After the first hour of the reaction, approximately one-third of PrK bound to the substrate, with no major differences observed between the samples (Figure 3A-B). However, from the 2-hour mark onward, significant differences emerged among the three conditions on _V_PLA (Figure 3A). When PrK was used alone, its bound molar fraction (χ_*PrK*_) remained relatively stable within the first 6 hours, reaching a final bound fraction of 52%. In contrast, the addition of Lip significantly enhanced PrK binding to the polymer, with the bound fraction steadily increasing throughout the reaction, reaching 85% after 20 hours. The addition of LOX I-B resulted in an intermediate effect. While it promoted PrK binding to the substrate, the increase was less pronounced compared to Lip, leading to an χ_*PrK*_ increase of approximately 33% more PrK bound to the polymer over the course of the reaction.

On _PC_PLA, similar results were observed with the key difference being a reduction in the χ_*PròK*_ bound to the substrate in the presence of LOX I-B compared to that of PrK alone. The overall effect of LOX I-B was reduced to only a 10% increase in the χ_*PrK*_. This finding aligns with the observed reduced synergistic effect mediated by LOX I-B on _PC_PLA in terms of LA release (Figure 2A). Overall, these results indicate that Lip, and to a lesser extent LOX I-B, enhanced the binding efficiency of PrK. This effect is likely due to an increase in the number of BS available for PrK on the substrate, ultimately leading to a higher depolymerization yield.

To further investigate the effect of Lip and LOX I-B on the polymer, a sequential enzyme addition was performed (Figure 3C). In this setup, the _V_PLA film was first incubated with an enzyme (E1) and after 24 hours a second enzyme was added (E2) keeping the _V_PLA film inside the test tubes. The data reported in Figure 3D are relative to the lactic acid measured at 48h time-point instrumental to observe if one of the enzyme combination could contribute to the hydrolysis during the course of the reaction of PrK. To our surprise none of the enzymatic pre-treatments with Lip or LOX for 24h led to an amelioration of the catalytic performance of the PrK in the successive 24h. Overall, 1.3 and 1.2 g/L of lactic acid were measured for the Lip and Lox pretreated _V_PLA, while 1.2 g/L was obtained for the sample just hydrolysed with PrK (without enzymatic pretreatment). Furthermore, an addition of Lip or Lox after an initial pre-digestion of the _V_PLA film with PrK did not synergise; instead it led to an inhibition of the latter enzymes. During the second 24 hours of digestion the transformation of _V_PLA into lactic acid reached 12.1 and 11.4 % instead of the 17.1 % obtained by the sole PrK.

## Discussion

The present study investigated the synergistic interaction between Proteinase K (PrK), a well-established enzyme with PLA-degrading activity, and two commercial enzymes, either hydrolytic, *Candida rugosa* lipase (CrLip) or redox, *Glycine max* lipoxygenase (LOX I-B). Lip and LOX I-B did not generate a detectable release of monomers or soluble oligomers when incubate with two kinds of PLA (lab-casted and postconsumer); however, both enzymes significantly boosted the PrK-mediated PLA degradation.

Lip enhanced PrK-mediated LA release by 5.7-fold on solvent-cast _V_PLA films and by 2.7-fold on _PC_PLA films. Additionally, the presence of calcium further increased the overall hydrolysis yield, reaching 43.5% for _V_PLA and 31% for _PC_PLA within 24 hours, in the presence of Lip and PrK. This increase in PLA hydrolysis aligns with previous reports indicating that PrK exhibits reduced thermal stability [26] and activity [22] in the absence of Ca^2+^. However, PLA hydrolysis plateaued after 48 hours, likely due to the thermal inactivation of PrK and its autoproteolytic behaviour, which becomes more pronounced at low enzyme concentrations [27]. This performance markedly exceeds that reported by Sourkouni et al. (2023) [28], who achieved only ∼40% PLA weight loss after 6 weeks using a comparable dosage of a complex enzymatic cocktail composed of five enzymes, including both commercial lipases and proteases. In contrast, our system achieved approximately 60% degradation of _V_PLA within only 72 hours. This highlights the superior efficiency of our approach, particularly when supplemented with Ca^2^□, in accelerating PLA depolymerization under mild conditions.

Lip appears to enhance PrK activity through two potential mechanisms: (i) increasing the number of productive binding sites for PrK on the polymer and perhaps (ii) hydrolysing soluble oligomers released by PrK activity. The first hypothesis is supported by the observation that, in the presence of Lip, the molar fraction of PrK bound to the polymer increased by an average of 70% on _V_PLA and 41% on _PC_PLA compared to PrK alone. This suggests that Lip may function as a PLA-modifying rather than a PLA-degrading enzyme [29], altering the polymer surface to enhance PrK binding. This hypothesis is further reinforced by previous studies showing that while Lip did not release LA from PLA [21], it could still induce surface modifications, such as small cracks in nonwoven PLA fibres [19, 30]. However, if added to the same previous setting but still having present the _V_PLA film, the Lip did not synergise with PrK. It appears that the synergistic action between these two hydrolytic enzymes lies in a simultaneous concerted action over the polymer chains, so as to promote the PrK binding to the PLA film and removal of the generated PLA-oligomers.

Similarly, LOX I-B exerted a significant synergistic effect when combined with PrK, resulting in a 2.5-fold increase in _V_PLA hydrolysis and 57% in _PC_PLA when used at a similar dosage to Lip. The observed synergism can be attributed to the increased χ_*PrK*_ bound to the substrate in the presence of LOX I-B, although not to the same extent as Lip. This effect was particularly pronounced for _V_PLA, where LOX I-B led to a greater enhancement of the hydrolysis yield. This suggests that LOX I-B also enhanced PrK activity by facilitating its adsorption onto the polymer surface. Moreover, in support of the hypothesis that LOX I-B plays a catalytic role in the depolymerization of PLA, the removal of molecular oxygen resulted in a significant loss of synergism with PrK. This O_2_-dependence provides further evidence that LOX I-B actively participates in the catalysis of PLA, and small amount of pyruvate was detected as a result of their incubations with _V_PLA. However, LOX I-B is a 13-lipoxygenase, which specifically oxidises the C13 position of linoleic acid within cis,cis-1,4-pentadiene moieties, which are lacking in PLA. Nevertheless, this represents only the primary reaction catalysed in vivo, as LOXs are also known to facilitate the subsequent conversion of hydroperoxides into various secondary oxidation products [31] and to mediate nonspecific oxidative reactions. For instance, Enoki et al. (2003) [32] reported a reduction in the molecular weight of trans-1,4-polyisoprene following incubation with LOX and its oxidative mediator, linoleic acid. More recently, Giannì et al. (2018) [33] documented LOX-mediated oxidation and fragmentation of lignin via oxygen- and carbon-centred radicals, both in the presence and absence of linoleic acid working as mediator. Given the known oxidative capabilities of LOX I-B, its potential activity toward PLA may involve the initiation of radical-mediated reactions. LOX I-B could facilitate the formation of reactive oxygen species or radical intermediates on the PLA-oligomers, perhaps, which may, in turn, interact with the polymer backbone. This could lead to oxidative modifications, such as chain scission or crosslinking, depending on the reaction conditions and the nature of the formed radicals. However, this oxidative activity could also negatively impact PrK-mediated hydrolysis. In the sequential enzyme addition experiment, PrK activity was impaired when LOX I-B was introduced last, suggesting that oxidative modifications of the substrate might hinder subsequent enzymatic hydrolysis.

This study did not aim to optimise enzyme ratios or reaction conditions for maximal PLA degradation; however, it demonstrated that the combined action of PrK and Lip or LOX I-B, despite their low molar concentration (i.e., approximately 2% of that of PrK) substantially improved PLA degradation. This finding is particularly relevant for future research, as leveraging such synergism could allow for a reduction in the overall enzyme loading required for effective PLA depolymerization. By enhancing degradation efficiency with minimal additional catalyst input, this approach represents a promising step toward a more cost-effective and scalable enzymatic recycling strategy for plastic waste [34].

